# Fenchel duality of Cox partial likelihood and its application in survival kernel learning

**DOI:** 10.1101/2020.05.04.077263

**Authors:** Christopher M. Wilson, Kaiqiao Li, Qiang Sun, Pei Fen Kuan, Xuefeng Wang

## Abstract

The Cox proportional hazard model is the most widely used method in modeling time-to-event data in the health sciences. A common form of the loss function in machine learning for survival data is also mainly based on Cox partial likelihood function, due to its simplicity. However, the optimization problem becomes intractable when more complicated regularization is employed with the Cox loss function. In this paper, we show that a convex conjugate function of Cox loss function based on Fenchel Duality exists, and this provides an alternative framework to optimization based on the primal form. Furthermore, the dual form suggests an efficient algorithm for solving the kernel learning problem with censored survival outcomes. We illustrate the application of the derived duality form of Cox partial likelihood loss in the multiple kernel learning setting

## 1. Introduction

The two most widely utilized models for censored survival data are the Cox proportional hazard (PH) model [1] and accelerated failure time (AFT), due to their flexibility and efficiency [2]. Cox PH models are the most widely used model in health and clinical sciences, while the AFT model is more popular in areas like engineering. The Cox model is a semi-parametric regression method where no assumptions imposed on the baseline hazard function. The parametric regression coefficients quantify the effect size of each covariate and the exponential of the coefficient is interpreted as the unit increase of the hazard ratio. The Cox model works well in practice because it can tolerate a modest deviation from the PH assumption. The partial likelihood in the Cox model is defined as the probability that one individual will experience the event at a time *t* among those who have survived longer than *t*. It was shown that maximizing partial likelihood provides an asymptotically efficient estimation of regression coefficients [3]. The Cox model has been successfully extended to various high-dimensional settings, where the number of features is more than the number of samples. Additionally, the log partial likelihood (LPL) is differentiable and convex. Thus the combination of Cox LPL, and l1 (lasso), l2 (ridge), or elastic net penalties can be directly solved by standard Newton-Rapshon method. Li and Luan [4] pioneered methods for kernel Cox regression in the framework the penalization framework from the view of function estimation in reproducing kernel Hilbert spaces. Furthermore, the convex combined loss function often guarantees the convergence of efficient optimization algorithms including coordinate descent [5], which is the core algorithm implemented in a popular R package for penalized regression **glmnet**. The Cox LPL has also been adopted in many machine learning approaches as a loss function to the survival setting. For example, Ridgeway [6] adapted the gradient boosting method for the Cox model, which is implemented in the R package **gbm**. Li and Luan [7] also considered a boosting procedure using smoothing splines to estimate the proportional hazards models. More recently, the application of neural network-based deep learning techniques to the Cox PH model has begun to receive attention [8]. These machine learning techniques generalize the Cox model to include non-linear effects and to better address heterogeneous effects.

In this paper, we derive the Fenchel dual form of Cox partial likelihood, which is a key step in the implementation of machine learning approaches for survival outcomes that can incorporate multiple high throughput data sources. Duality approach is a basic tool in machine learning and is commonly used in nonlinear programming and convex optimization to provide a lower bound approximation for the primal problem. It is often easier to optimize the lower bound via the dual form and it has fewer variables in the high dimensional setting. The main property of the resulting function, called Fenchel conjugate, is always convex regardless of the convexity of the original function. Note that the Lagrangian and Fenchel dual are defined under different contexts, even though many Lagrangian duals can be derived from Fenchel conjugate functions and in many cases, both of them are referred to as a “dual problem”. Lagrangian duality form is defined within the context of the optimization problem (often with constraints), while the Fenchel form is more general and is defined for a function. Our work is motivated by the need to bridge the gap between modern machine learning techniques and survival models. To apply methods like SVM, survival data are often dichotomized with a cutoff time point. Such a method will yield biased results because censored data points are excluded from the analysis, additionally, the results will be affected by different cutoff values.

The remainder of the paper is organized as follows. In Section 2, we review the Cox proportional hazard model and Fenchel duality. In Section 3, we derive a conjugate function for the Cox model. We perform simulations in Section 4 to demonstrate the usage of the derived form in the multiple kernel learning. We analyze both Skin Cutaneous Melanoma (SKCM) gene and miRNA expression data from The Cancer Genome Atlas (TCGA). Finally we conclude with a discussion in Section 6.

## 2. Methods

### 2.1. Fenchel duality

Suppose we have function *f* (*x*) on *R*^*n*^, then the Fenchel convex conjugate of *f* (*x*) is defined in terms of the supremum by

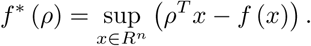

The mapping from *f* (.) to *f*∗(.) defined above is also known as the Legendre-Fenchel transform. The convex conjugate function measures the maximum gap between line function *ρ*^*T*^*x* and original function *f* (*x*), where each pair of *ρ*, *f*^*T*^ (*ρ*) corresponds to a tangent line of the original function *f* (*x*). The resulting function *f*∗ has the nice property to be always convex, because it is the supremum of an affine function. Figure 1 illustrates how the conjugate dual for a classic example *f* (*x*) = |*x*|. The conjugate function offers one important option to build a dual problem that might be more tractable or computationally efficient than the primal problem. Note in Figure 1A, when |*ρ*| > 1 that as *x* → ∞ then |*ρ*|*x* → ∞, thus sup_*x*_{*ρx* − |*x*|} = ∞. Alternative, when |*ρ*| ≤ 1 that |*ρ*|*x* ≤ *x* for all *x*, hence the largest value for sup_*x*_{*ρx* − |*x*|} = 0, when *x* = 0. The complex conjugate is illustrated in Figure 1B, note that is a convex function.

**Figure 1:**
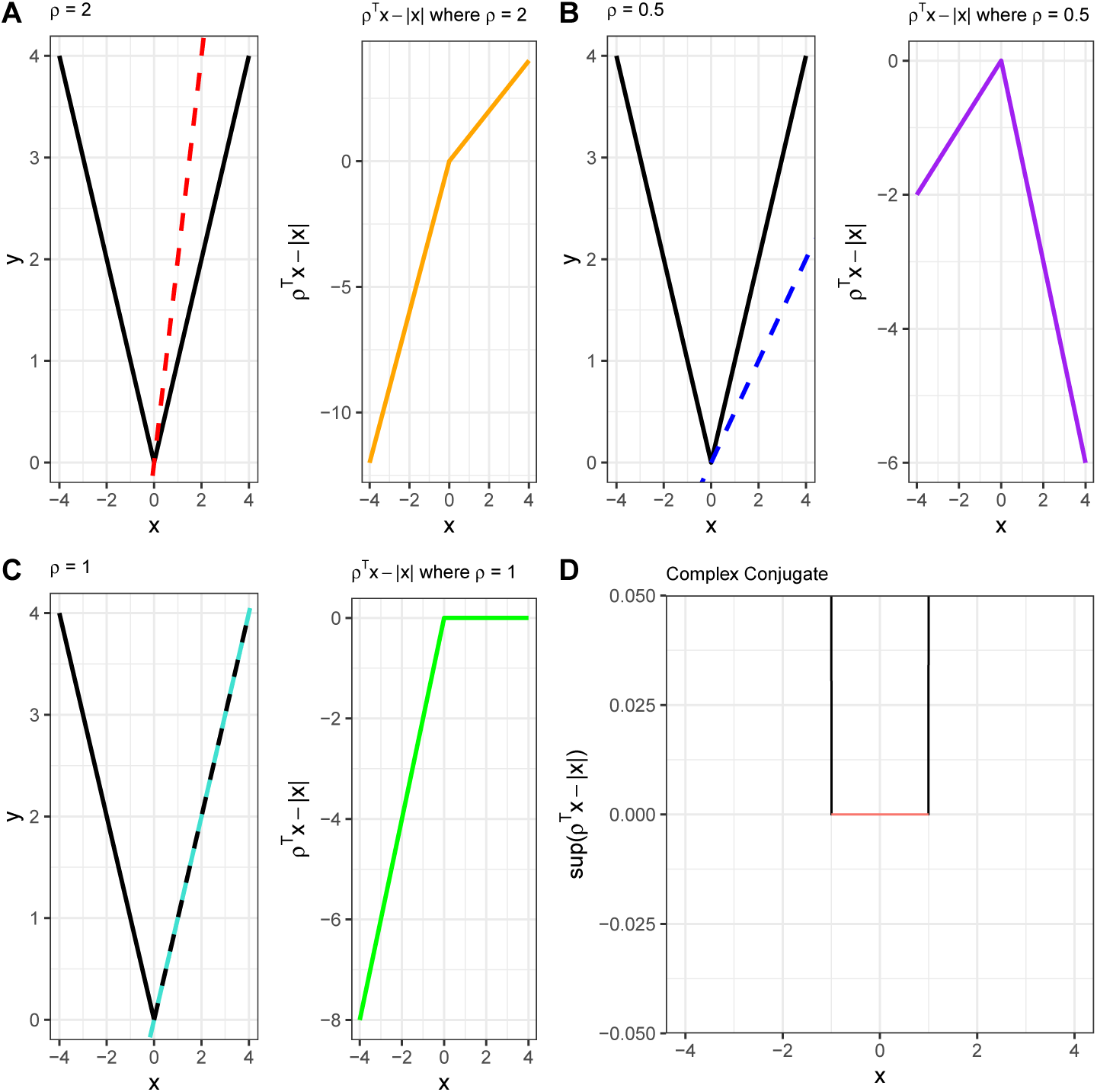
“Visual derivation” of the complex conjugate function is computed. (A)-(C) Shows the properties of *ρx* − *f* (*x*), where *f* (*x*) = |*x*| for particular values for *ρ* = 2, 1, 0.5. Notice that the difference between *ρx* and *f* (*x*) remains less than infinity only when |*ρ*| ≤ 1, while when *ρ* > 0, *ρx* increases faster than *f* (*x*). The final form of the conplex conjugate is displayed in (D).

By Fenchel-Moreau theorem, *f* = *f*∗∗ if only and only if *f* is a convex and and lower-semi continuous function which holds for Cox proportional hazards model. Therefore, we can convert the problem into dual problem with *f*∗ to obtain 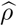 and map it back to our primal and obtain final solution. We define the relative interior of a set *C* as *ri*(*C*) = {*x* ∈ *C*| for all *y* ∈ *C* there exists *λ* > 1 such that *λx* + (1 − *λ*)*y* ∈ *C*}, in other words for an point *x* ∈ *C* there exists a ball that is entirety contained in *C*. Additionally, using Fenchel duality theorem[9] (Theorem 31.1), we have the following statement. If *ri* (*dom* (*f*)) ∩ *ri* (*dom* (*g*)) ≠ ∅, and *f* (⋅) and *g* (⋅) are convex, we have

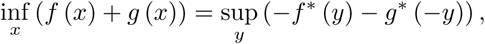

note that minimizing *h*(*x*) = *f* (*x*) + *g*(*x*) occurs when ∇*h* = ∇*f* + ∇*g* = 0 or when ∇*f* = −∇*g*. Hence, Fenchel duality theorem shows us that minimizing the summation of two convex function can be reduced to the problem of maximizing the gap between their parallel tangent lines since it is the lower bound of primary problem. In our case, both the Cox loss function and EN regularizer are both convex, so we can apply this theorem that our target problem which reduces to the following problem

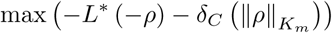

where *L*∗ and *ϕ* are the conjugate function of *L*, *K*_*m*_ Hilbert space that is generated by the reporducing kernel *K*_*m*_, and *ϕ*.

### 2.2. Cox proportional hazard model and partial likelihood

The Cox PH model [1] relates the covariates to the hazard function of the outcome at time *t* using the following equation,

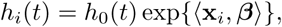

where *h*_0_(*t*) is the baseline hazard function at time *t* and **x**_*i*_ is the vector of predictor variables for the *i*^*th*^ subject. An appealing feature of the Cox model is that, as shown in the partial likelihood

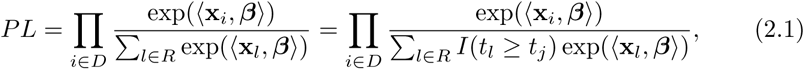

estimates of regression coefficients are obtained without parametric assumptions about the baseline hazard function. Here *D* is the set of uncensored subjects and *R* is the set of the observations at risk at time *t*. The PL can be understood as constructing the conditional probability that the event occurs to a particular subject at time *t*. Typically, we optimize the negative log of Cox PL 2.1,

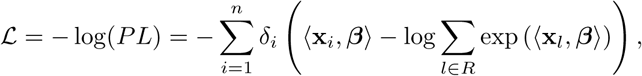

where *δ* is the event indicator. It can be shown that *𝓛* is convex and thus obtaining the regression coefficients that minimize *𝓛* can be conducted by using gradient based methods. This framework can be extended to penalized regression by adding a regularization term to *𝓛*. For example, the lasso solution of regression coefficients corresponds to 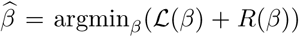, where *R*(*β*) denotes the regularization terms applied to constrain coefficients, such as the l1/l2 or group lasso penalty terms [10].

### 2.3. Multiple Kernel Learning

In the traditional Cox regression framework, we assume a linear relationship between our predictors and survival time. However, in reality, the relationship is far more complex. Meanwhile, nonlinear models are often hard to analyze and interpret. Kernel methods are non-parametric methods that utilize reproducing kernel Hilbert space (RKHS) [11, 12], and provide a useful alternative to linear or nonlinear models. Kernel functions map a predictor matrix from *n*×*p* to *n*×*n*. This allows us to focus on the similarity between subjects dramatically reducing the complexity of the predictor space to finding a linear relationship between a similarity measure. For instance, using polynomial kernel, we can map a circle boundary problem to a linear boundary problem, which dramatically reduces our computation cost.

The exact relationships between predictions and outcomes are unknown, hence selecting an optimal kernel function presents a challenge. There are no clear rules for selecting a single optimal kernel, but cross-validation is usually implemented[13, 14, 15]. An interesting property of kernels is that a linear combination of two kernel functions results in another kernel function [11]. There have been many works that utilize this fact and have shown that convex combinations of multiple kernels can provide more accurate classifiers than single kernels[13, 16, 17]. Learning the optimal kernel weights is referred to as multiple kernel learning (MKL).

Under the MKL framework, we can denote our target problem as follow

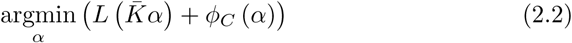

where *L* (⋅) is the loss function of Cox proportional hazards model, 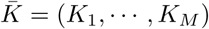 are a set of kernels, *α* is the coefficient matrix for each kernels, and *ϕ*_*C*_ (⋅) is the regularized term for coefficients matrix. In this paper we used elastic net (EN) penalty which can be written as

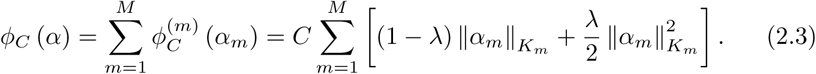

where *λ* ∈ [0, 1] defines the amount of weight is assigned to the *l*2 and *l*1 regularizer, and *C* is a multiplier that changes the impact prediction error on the objective function in 2.2.The EN penalty allows us strike a balance between the *l*1 and *l*2 regularization. In other words, it allows for sparse selection that the coefficients for non-informative kernels will be shrunk to zeros, and the coefficients for similar kernels tend to be close, so called group property, which returns us a consistent result [18, 17]. The elastic net penalty is non-smooth, we can use the theory of Moreau’s envelope function obtain the approach to solve this problem.

### 2.4. Moreau Envelope and Elastic Net

The Moreau envelope function allows us to approximate a non-smooth function with a smooth function leading to simpler optimization task. Let 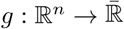 be a function, for every *γ* > 0 we define the Moreau envelope as

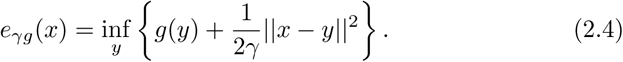

The Moreau envelope strikes a balance between function approximation and smoothness through the parameter *γ*. Note that Moreau envelope of *g*(*y*) = *−ρ*^*T*^*y* is the negative of the convex conjugate of 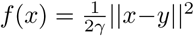. Additionally, if *e*_*γg*_ is smooth derivative given by

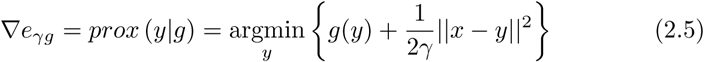

A simple example of the Moreau envelope is shown in Figure 2A. Note that as *γ* increases the approximation of the function becomes worse and the shape of the function around *x* = 0 is increasingly rounded.

**Figure 2:**
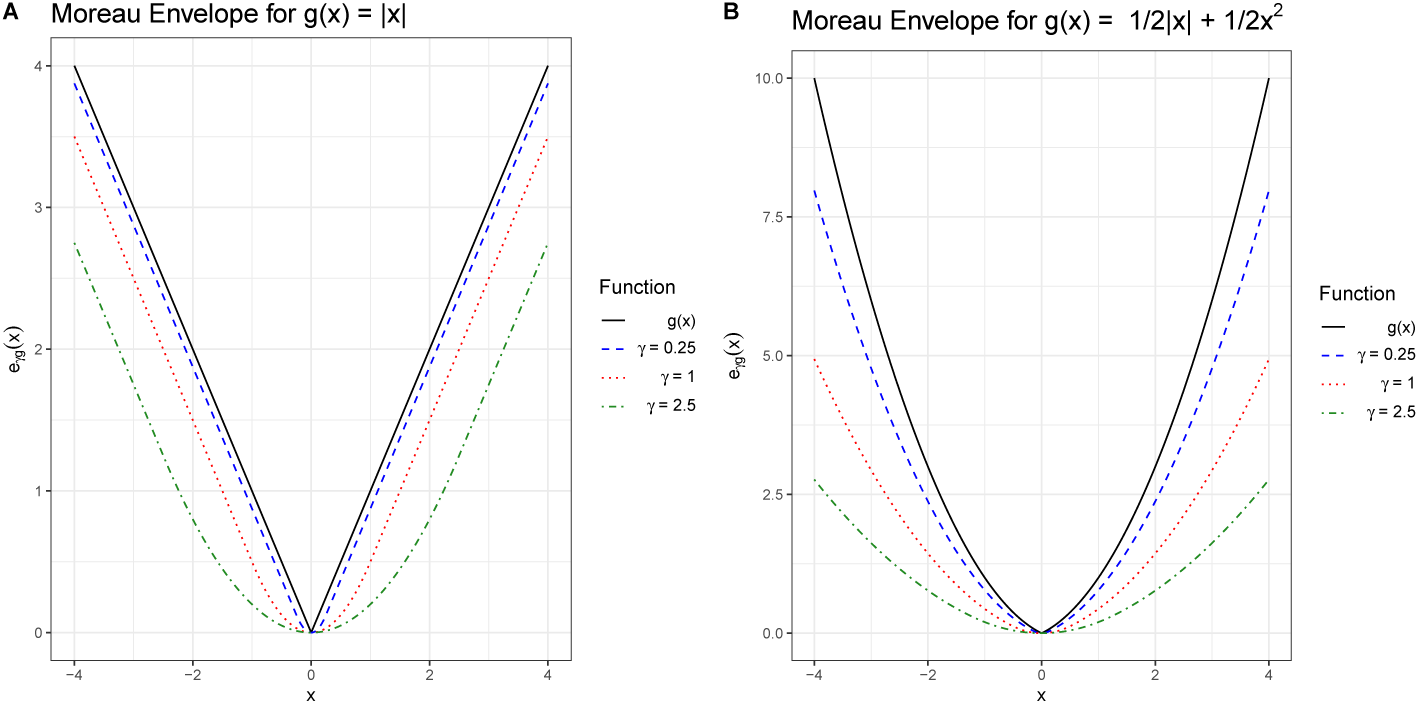
Moreau envelope for lasso regularizer *g*(*x*) = |*x*| for several values of smoothing parameter γ, and elastic net regularizer *g*(*x*) = 1/2|*x*| + 1/2*x*^2^. Notice that as the value for *γ* increases we obtain a function that is more smooth, but a worse approximation of *g*(*x*).

Now we apply the concept of the Moreau envelope to the EN. We set *x* = *α*_*m*_, *y* = *α′*_*m*_, and *γ* = 1 from 2.5 resulting in

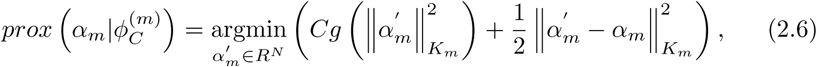

where 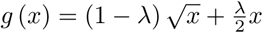.

Using Cauchy-Schwarz inequality we have

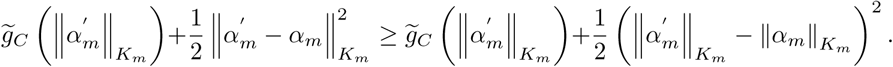

Therefore, we can obtain the minimum solution that

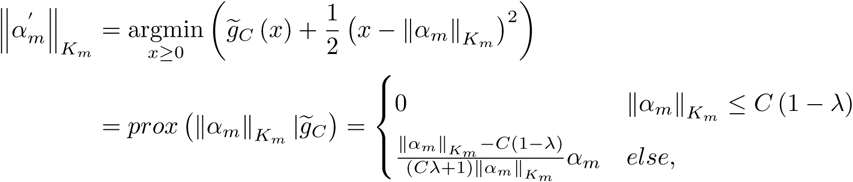

where *prox* (⋅) is known as the soft operator, see 2B.

## 3. SpicyMKL algorithm for Cox Proportional Hazard

SpicyMKL was introduced as an efficient implementation of MKL what could learn the best convex combination of potentially 1000 candidate kernels. In order to accomplish this the MKL problem was reformulated using the Moreau envelope and the convex conjugate to ensure that the sum of the loss and regularization penalties is a smooth and convex function [17]. SpicyMKL was introduced for multiple loss function and regularization function, but was not extended to the survival setting.

In our problem, we denote *L*∗ (⋅) as the conjugate function of the Cox loss function

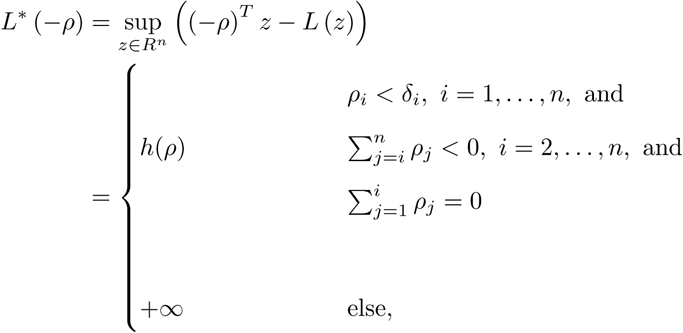

where

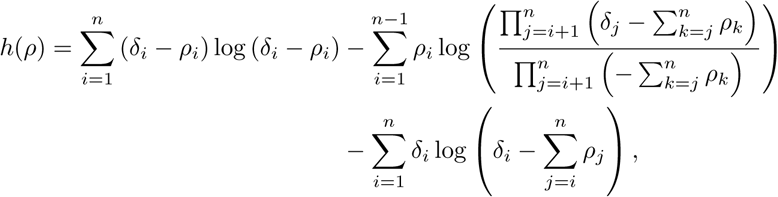

and 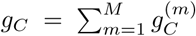 is the conjugate function of the soft operator of the regularizer,

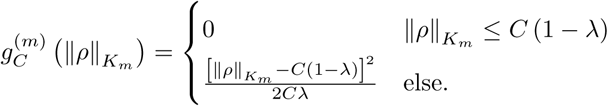

A full derivation of the results can be found in the supplemental materials. We can see that both *L*∗ and *δ*_*C*_ are secondarily differentiable, we can use Newton method to solve above questions.

The conjugate function of *L* is secondarily differentiable where the first derivative is given by

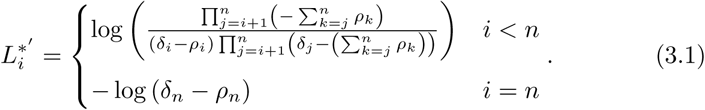

We can see that if *j* < *i*, the derivation respected to the *i*^*th*^ component does not contain *j*, so the 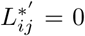. If *j* > *i*, 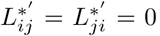. Hence the Hessian matrix of the conjugate function is diagonal matrix, then we can obtain by taking the second derivative,

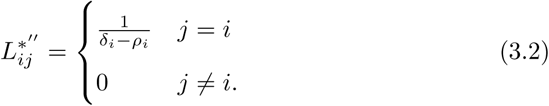

We can calculate gradient and Hessian matrix of 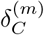 using (3.1) and (3.2) which are given by

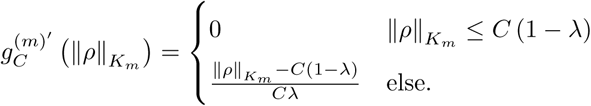

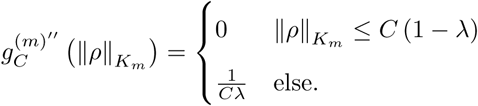

We can see that the conjugate function is feasible if *λ* > 0, which means we can only use the smooth dual form for elastic net but not block one norm penalty. The block one norm penalty is a kernelized version of group lasso [19, 20].

From derivation of the SpicyMKL algorithm, we have

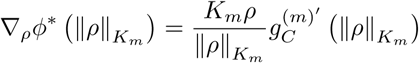

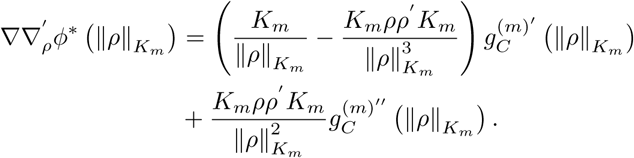

Using Newton algorithm we can obtain the optimal 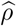. As shown in the supplemental material, the step size of Newton update is given by the size that will not make the update of *ρ* goes beyond the domain of *L*∗. To satisfy the 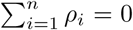 constraint for *L*∗, we added a penalty function 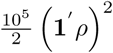. Using Rockafellar[9] (Theorem 31.3), we have

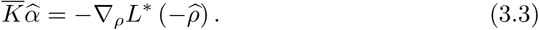

Under Karush–Kuhn–Tucker (KKT) condition.

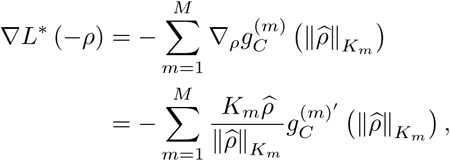

and the solution to (3.3) is

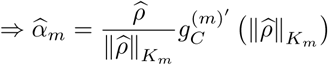

## 4. Simulation Data

To evaluate the performance of Multiple Kernel Cox regression (MKCox), we simulate data that are generated with different relationships between the feature and the hazard function. Our simulations were inspired by Katzman [8]. We simplify these simulations we conducted to benchmark and explore the properties of MKCox. We simulate the features from a bivariate normal distribution, **X** = [*X*_1_*, X*_2_], with *µ* = 0, and 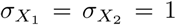 and a range of values of correlation, and we consider the following hazard function:

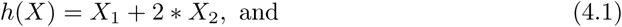

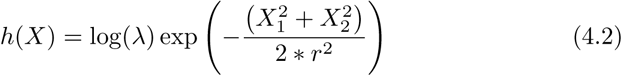

where *λ* = 5, and *r* = 1/2. Then the survival times are generated by:

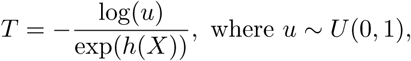

a censoring time is established, so that approximately 50% of patients have observed an event. We used two kernels in for our illustration *K*_1_ is the radial basis function with hyperparameter *σ* = 2, and *K*_2_ is a linear kernel. We compare MKCox to the following methods random survival forest ([21]) (RF) implemented in *randomForestSRC* (2.9.0), and stochastic gradient boosted ([22]) Cox regression (GBM) implemented in *gbm* (2.1.5). All our analyses were performed in R (3.6.0).

We aim to obtain a good estimate of the hazard function, as well as, maintain a good prediction of survival time. In Figures 3 and 4, we see that all methods can capture the structure of the hazard function when the underlying relationship is linear. Additionally, MKCox can recover the underlying patterns in the hazard function better than Cox regression and GBM, while both RSF and MKCox both capture the nonlinear pattern for accurately. To evaluate the performance of MKCox will compute popular metrics for survival models such as the concordance index. These results are shown in Table 1. Notice in the linear case that all methods provide similar concordance indices, while in the nonlinear case Cox performs substantially worse and MKCox slightly better than other machine learning methods. These simulations illustrate that MKCox can produce similar or better results under different underlying relationships between the hazard ratio and the features.

**Table 1:**
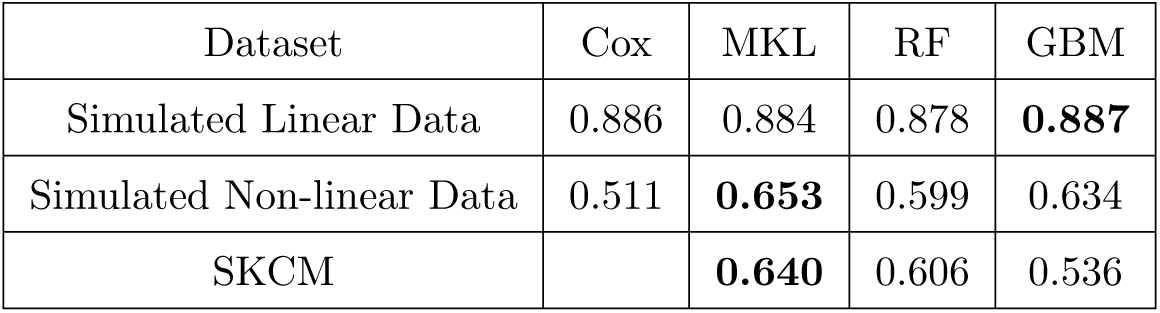
Concordance Index for Each Model

| Dataset | Cox | MKL | RF | GBM |
| --- | --- | --- | --- | --- |
| Simulated Linear Data | 0.886 | 0.884 | 0.878 | **0.887** |
| Simulated Non-linear Data | 0.511 | **0.653** | 0.599 | 0.634 |
| SKCM |  | **0.640** | 0.606 | 0.536 |

**Figure 3:**
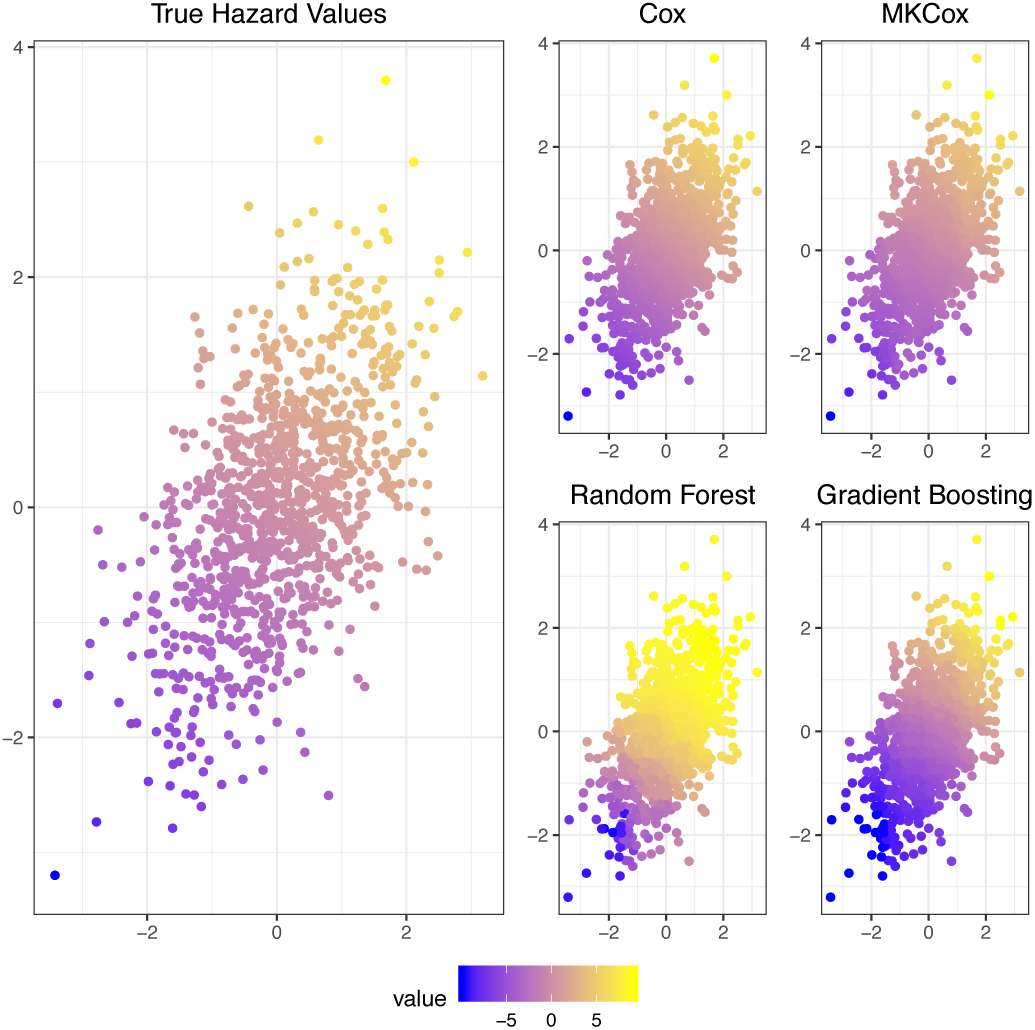
((A) The values of the linear hazard function used in (4.1), *h*(*x*_1_, *x*_2_) = *x*_1_ + 2*x*_2_. (B) Predicted hazard values by Cox proportional hazard, and the three machine learning techniques.

**Figure 4:**
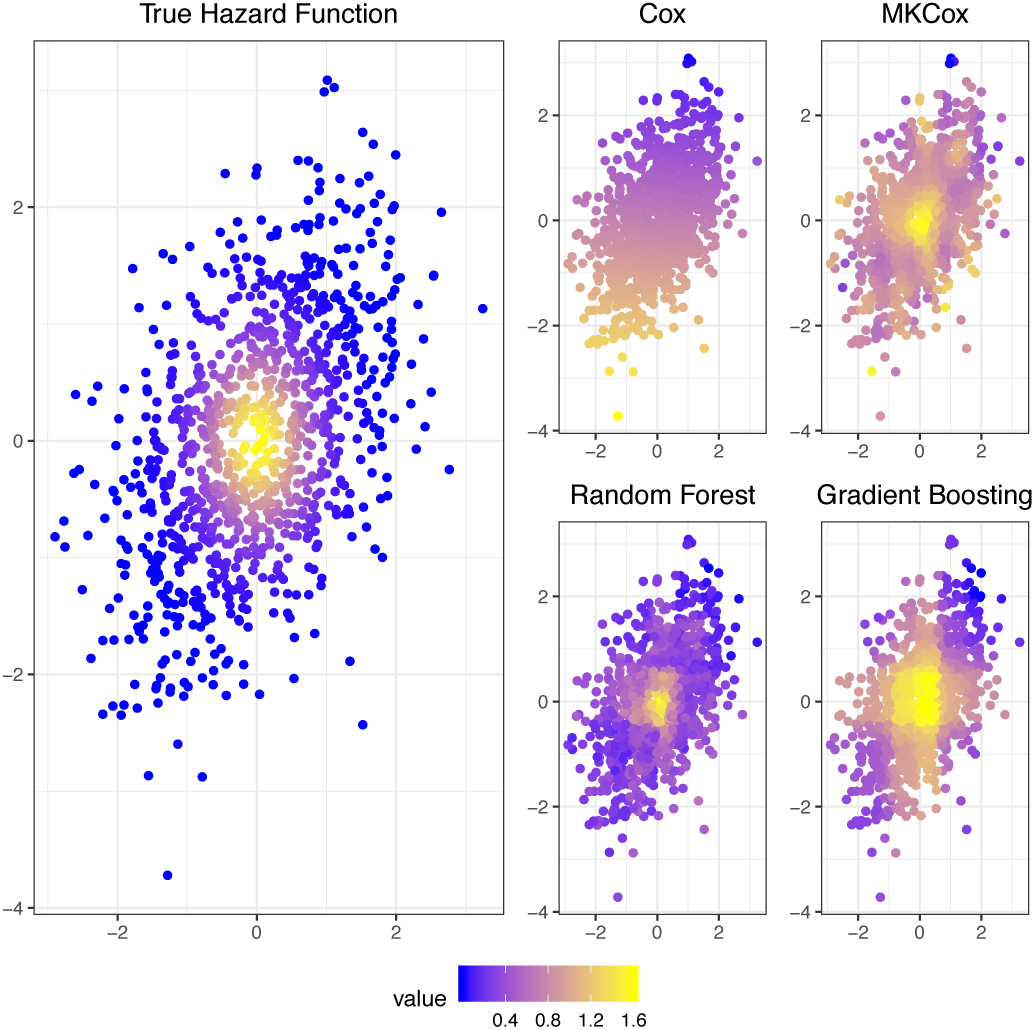
IA) The values of the nonlinear hazard function used in (4.1), 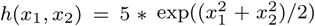. (B) Predicted hazard values by Cox proportional hazard, and the three machine learning techniques.

## 5. Case Study

The Cancer Genome Atlas (TCGA) project is a large initiative to study the multiomics effect of gene expression RNAseq and stem loop expression on patients’ survival time[23, 24]. We downloaded the data from Genomic Data Commons (GDC) Data Portal. The latest survival data were downloaded using the *TCGAbiolinks* ([25]) package in R. The gene expression and miRNA expression data were downloaded from University of California at Santa Cruz (UCSC) Xena ([26]) (https://xena.ucsc.edu/) database. For gene expression, we used the fragments per kilobase of transcript per million mapped reads upper quartile (FPKM-UQ) with log2 (*x* + 1) transformation on mRNA via high-throughput sequencing (HTseq) ([27]) technique with gencode v22, while for stem loop expression, we used the per million mapped reads (RPM) with log2 (*x* + 1) transformation via miRNA expression quantification technique aligned to GRCh38. In total, we have 235 dead and 215 alive (450 in total) patients.

Since we have two features sources (gene expression and stem loop expression), we kernelized the two feature sources separately using pathway kernels with 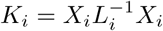 where *K*_*i*_, *X*_*i*_ and *L*_*i*_ are the kernel matrix, feature matrix and standardized Laplacian matrix for each feature source, respectively. So our model can be written as

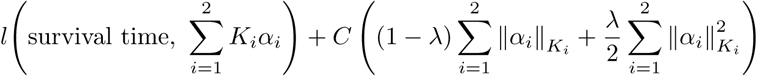

which is equivalent to a linear grouped network regularized model

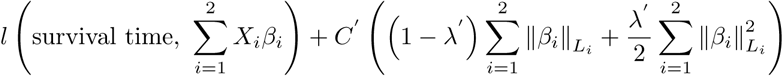

where 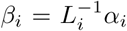 so that we can obtain the coefficient of feature *i* using this transformation. In this case the kernel learning method has strong interpretability. The Laplacian matrices were estimated empirically by neighbor network and coexpression network method proposed by [28].

To evaluate the performance we split that data into 301 training and 149 test samples, stratified by survival status (dead versus alive). The models we compared were all trained on training data and the results were obtained on test data. The parameters for our MKL model and GBM models were tuned by 5fold cross-validation. From Table 1 we can see that our proposed multiple kernel learning using network kernels worked the best. Though it was a linear model, it achieved a higher concordance index than nonlinear tree-based models like the random forest or stochastic gradient boosting machine. Due to the flexibility and efficiency of MKCox can incorporate many pathways under different kernel representations.

## 6. Conclusion

In this paper, we derived an efficient multiple kernel learning algorithm for survival prediction models and the convex conjugate function for Cox proportional hazards loss function. A challenge of deriving efficient algorithms for proportional hazard models is that the Hessian is not a diagonal matrix. However, through the convex conjugate function we can utilize the diagonal property to achieve a time competitive algorithm. Therefore, the Cox proportional hazards loss function can be more easily implemented than other machine learning methods.

Both the simulation and case study in our paper showed a robust performance of our proposed method in likelihood function estimation and out of data prediction, even compared to tree-based methods which often showed a strong predictive power. As we can see that MKL method was shown to have superior performance in cancer genomic studies ([29]). Future studies include extending the model to other more complex survival problems including competing risks.

## Supporting information

Derivations

## 7. Acknowledgements

This work was supported in part by Institutional Research Grant number 14-189-19 from the American Cancer Society, and Moffitt Foundation Methodological Development Pilot Grant.

