## Supplementary material for "Fenchel duality of Cox partial likelihood and its application in survival kernel learning": Derivations

### 1. Supplemental Material

#### *1.1. Conjugate function*

We provide a detailed derivation for conjugate function from the Cox partial likelihood. The conjugate function is given by

$$\begin{aligned} L^*(-\rho) &= \sup_z \left( (-\rho)' z - L(z) \right) \\ &= \sup_z \left( \sum_{i=1}^{n} \left[ -(\rho_i - \delta_i) z_i - \delta_i \log \sum_{j=i}^{n} \exp(z_j) \right] \right), \end{aligned}$$

where $z = \mathbf{X}'\boldsymbol{\beta}$ and $\delta \in \{0,1\}^n$ indicating whether an event occurred. We also assume that the time to events or censoring times are sorted in ascending order, $\min(t_1, C_1) < \min(t_2, C_2) < \cdots < \min(t_n, C_n)$ and assuming that there are no ties. This the risk set at $t_i$ is subjects $\{(i+1), \ldots, n\}$.

Let

$$g = \sum_{i=1}^{n} \left( -(\rho_i - \delta_i) z_i - \delta_i \log \sum_{j=i}^{n} \exp(z_j) \right), \tag{1.1}$$

we take the partial derivative of $g(\cdot)$ with respect to $z_i$ and set them to be zero,

$$\nabla g_i = \frac{\partial g}{\partial z_i} = -\rho_i + \delta_i - \exp(z_i) \sum_{j=1}^{i} \delta_j \left( \sum_{k=j}^{n} \exp(z_k) \right)^{-1} = 0,$$

note that $\exp(z_i)(\sum_{k=j}^{n} \exp(z_k))^{-1} > 0$ and $\delta_i \exp(z_i)(\sum_{k=j}^{n} \exp(z_k))^{-1} \geq 0$, hence $\delta_i - \rho_i \geq 0$.

This leads to the following condition:

$$0 \leq \delta_i - \rho_i = \exp(z_i) \sum_{j=1}^{i} \delta_j \left( \sum_{k=j}^{n} \exp(z_k) \right)^{-1} \leq \sum_{j=1}^{i} \delta_j. \tag{1.2}$$

Let $h_i = \exp(z_i) > 0$, we have

$$\delta_1 - \rho_1 - \frac{\delta_1 h_1}{h_1 + \cdots + h_n} = 0 \qquad (1.3)$$

$$\delta_2 - \rho_2 - \frac{\delta_1 h_2}{h_1 + \cdots + h_n} - \frac{\delta_2 h_2}{h_2 + \cdots + h_n} = 0 \qquad (1.4)$$

$$\vdots$$

$$\delta_n - \rho_n - \frac{\delta_1 h_n}{h_1 + \cdots + h_n} - \frac{\delta_2 h_n}{h_2 + \cdots + h_n} - \cdots - \frac{\delta_n h_n}{h_n} = 0. \qquad (1.5)$$

Now we compute the sum of system of equations above which yields

$$\sum_{i=1}^{n} \rho_i = 0. \qquad (1.6)$$

Since we have

$$\delta_i - \rho_i - h_i \sum_{j=1}^{i} \delta_j \left( \sum_{k=j}^{n} h_k \right)^{-1} = 0, \text{ and} \qquad (1.7)$$

$$\delta_{i+1} - \rho_{i+1} - h_{i+1} \sum_{j=1}^{i+1} \delta_j \left( \sum_{k=j}^{n} h_k \right)^{-1} = 0, \qquad (1.8)$$

we can divide **??** by $h_i$ and **??** by $h_{i+1}$ and subtract to obtain

$$\frac{\delta_{i+1} - \rho_{i+1}}{h_{i+1}} - \frac{\delta_i - \rho_i}{h_i} - \frac{\delta_{i+1}}{\sum_{j=i+1}^{n} h_j} = 0. \qquad (1.9)$$

Solve **??** at $i = n - 1$, we obtain

$$\frac{h_n}{h_{n-1}} = \frac{-\rho_n}{\delta_{n-1} - \rho_{n-1}} = \frac{-\rho_n \left(\delta_n - \rho_n\right)}{\left(\delta_{n-1} - \rho_{n-1}\right)\left(\delta_n - \rho_n\right)}$$

$$\sum_{i=n-1}^{n} h_i = \frac{\delta_{n-1} - \rho_n - \rho_{n-1}}{\delta_{n-1} - \rho_{n-1}} h_{n-1}.$$

We can use mathematical induction to prove that

$$\frac{h_{i+1}}{h_i} = \frac{\left(\delta_{i+1} - \rho_{i+1}\right)\left(-\sum_{j=i+1}^{n} \rho_j\right)}{\left(\delta_i - \rho_i\right)\left[\delta_{i+1} - \left(\sum_{j=i+1}^{n} \rho_j\right)\right]}, \text{ and } \sum_{j=i}^{n} h_i = \frac{\delta_i - \left(\sum_{j=i}^{n} \rho_j\right)}{\delta_i - \rho_i} h_{i-1}$$

Suppose that for $i = k$, $1 < k < n$ we have the following equation hold, using equation **??** we have

$$
\begin{aligned}
&\frac{\delta_k-\rho_k}{h_k}-\frac{\delta_{k-1}-\rho_{k-1}}{h_{k-1}}-\frac{\delta_k\left(\delta_k-\rho_k\right)}{\left[\delta_k-\left(\sum_{j=k}^n\rho_j\right)\right]h_k}\\
&\qquad=\frac{\delta_k-\rho_k}{h_k}-\frac{\delta_{k-1}-\rho_{k-1}}{h_{k-1}}-\frac{\delta_k}{\sum_{j=i}^n h_i}=0,
\end{aligned}
$$

which implies that

$$
\frac{h_k}{h_{k-1}}=\frac{-\left(\delta_k-\rho_k\right)\sum_{j=k}^n\rho_j}{\left(\delta_{k-1}-\rho_{k-1}\right)\left[\delta_k-\left(\sum_{j=k}^n\rho_j\right)\right]},
$$

and

$$
\begin{aligned}
\sum_{j=k-1}^n h_j&=h_{k-1}+\sum_{j=k}^n h_k=h_{k-1}+\frac{\delta_k-\left(\sum_{j=k}^n\rho_j\right)}{\delta_k-\rho_k}\frac{-\left(\delta_k-\rho_k\right)\sum_{j=k}^n\rho_j}{\left(\delta_{k-1}-\rho_{k-1}\right)\left[\delta_k-\left(\sum_{j=k}^n\rho_j\right)\right]}h_{k-1}\\
&=\frac{\delta_{k-1}-\left(\sum_{j=k-1}^n\rho_j\right)}{\delta_{k-1}-\rho_{k-1}}h_{k-1}.
\end{aligned}
$$

Therefore we have for $i=1,\cdots,n$

$$
h_i=\left(\prod_{j=i}^{n-1}\frac{h_j}{h_{j+1}}\right)h_n=\frac{\left(\delta_i-\rho_i\right)\prod_{j=i+1}^n\left(\delta_j-\left(\sum_{k=j}^n\rho_k\right)\right)}{\left(\delta_n-\rho_n\right)\prod_{j=i+1}^n\left(-\sum_{k=j}^n\rho_k\right)}h_n \tag{1.10}
$$

$$
\sum_{j=i}^n h_j=\frac{\prod_{j=i}^n\left(\delta_j-\left(\sum_{k=j}^n\rho_k\right)\right)}{\left(\delta_n-\rho_n\right)\prod_{j=i+1}^n\left(-\sum_{k=j}^n\rho_k\right)}h_n. \tag{1.11}
$$

We can plug **??** and **??** into function $g\left(\cdot\right)$, **??**, and solve for $h_n$. In this way we can obtain the conjugate function

$$
\begin{aligned}
L^*=&\sum_{i=1}^{n-1}\left(\delta_i-\rho_i\right)\log\left(\frac{\left(\delta_i-\rho_i\right)\prod_{j=i+1}^n\left(\delta_j-\sum_{k=j}^n\rho_k\right)}{\left(\delta_n-\rho_n\right)\prod_{j=i+1}^n\left(-\sum_{k=j}^n\rho_k\right)}\;h_n\right)\\
&-\sum_{i=1}^{n-1}\delta_i\log\left(\frac{\prod_{j=i}^n\left(\delta_j-\left(\sum_{k=j}^n\rho_k\right)\right)}{\left(\delta_n-\rho_n\right)\prod_{j=i+1}^n\left(-\sum_{k=j}^n\rho_k\right)}\right)h_n+\left(\delta_n-\rho_n\right)\log\left(\frac{\delta_n-\rho_n}{\delta_n-\rho_n}h_n\right)-\delta_n\\
&=\sum_{i=1}^n\left(\delta_i-\rho_i\right)\log\left(\delta_i-\rho_i\right)+\log\left(\frac{\delta_n-\rho_n}{h_n}\right)\sum_{i=1}^n\rho_i\\
&-\sum_{i=1}^{n-1}\rho_i\log\left(\frac{\prod_{j=i+1}^n\left(\delta_j-\sum_{k=j}^n\rho_k\right)}{\prod_{j=i+1}^n\left(-\sum_{k=j}^n\rho_k\right)}\right)+\sum_{i=1}^{n-1}\delta_i\log\left(\frac{\prod_{j=i+1}^n\left(\delta_j-\sum_{k=j}^n\rho_k\right)}{\left(\delta_n-\rho_n\right)\prod_{j=i+1}^n\left(-\sum_{k=j}^n\rho_k\right)}\right)h_n
\end{aligned}
$$

$$-\sum_{i=1}^{n-1}\delta_i \log\left(\frac{\prod_{j=i}^{n}\left(\delta_j-\left(\sum_{k=j}^{n}\rho_k\right)\right)}{\left(\delta_n-\rho_n\right)\prod_{j=i+1}^{n}\left(-\sum_{k=j}^{n}\rho_k\right)}\right)h_n-\delta_i\log\left(\delta_n-\rho_n\right)$$

$$=\sum_{i=1}^{n}\left(\delta_i-\rho_i\right)\log\left(\delta_i-\rho_i\right)-\sum_{i=1}^{n-1}\rho_i\log\left(\frac{\prod_{j=i+1}^{n}\left(\delta_j-\sum_{k=j}^{n}\rho_k\right)}{\prod_{j=i+1}^{n}\left(-\sum_{k=j}^{n}\rho_k\right)}\right)$$

$$-\sum_{i=1}^{n}\delta_i\log\left(\delta_i-\sum_{j=i}^{n}\rho_i\right)+\log\frac{\delta_n-\rho_n}{h_n}\sum_{i=1}^{n}\rho_i$$

From **??**, the last term that includes $h_n$ is zero so the conjugate function reduces to

$$L^*(-\rho)=\sum_{i=1}^{n}\left(\delta_i-\rho_i\right)\log\left(\delta_i-\rho_i\right)-\sum_{i=1}^{n-1}\rho_i\log\left(\frac{\prod_{j=i+1}^{n}\left(\delta_j-\sum_{k=j}^{n}\rho_k\right)}{\prod_{j=i+1}^{n}\left(-\sum_{k=j}^{n}\rho_k\right)}\right)$$
$$-\sum_{i=1}^{n}\delta_i\log\left(\delta_i-\sum_{j=i}^{n}\rho_i\right)\qquad(1.12)$$

Note that if we take the second derivative of function $g(\cdot)$, **??**, the first term drops will zero, hence $\partial^2 g/\partial z_i^2$ is the same as Cox regression. Thus $g$ is a convex function. We can also write $L^*$ incorporating the constraints, **??**, **??**, and

$$L^*(-\rho)=\begin{cases}\sum_{i=1}^{n}\left(\delta_i-\rho_i\right)\log\left(\delta_i-\rho_i\right)-\sum_{i=1}^{n-1}\rho_i\log\left(\frac{\prod_{j=i+1}^{n}\left(\delta_j-\sum_{k=j}^{n}\rho_k\right)}{\prod_{j=i+1}^{n}\left(-\sum_{k=j}^{n}\rho_k\right)}\right) & 0<\rho_i<\delta_i, 1\le i\le n,\\ \qquad -\sum_{i=1}^{n}\delta_i\log\left(\delta_i-\sum_{j=i}^{n}\rho_i\right) & \sum_{j=i}^{n}\rho_j<0, 2\le i\le n, \text{ and}\\ & \sum_{j=1}^{i}\rho_j=0\\ +\infty & else\end{cases}$$

### 2. Step size of Newton update

In each iteration of Newton update, we have

$$\rho^{(t+1)}\leftarrow\rho^{(t)}-sH^{-1}\nabla,$$

where $H$ is the Hessian matrix, $\nabla$ is the gradient and $s$ is the step size. Let $\Delta=-H^{-1}g$ by apply the following constraints $\rho_i<\delta_i$ for $1\le i\le n$ and

$\sum_{j=i}^{n} \rho_j < 0$ for $2 \le i \le n$, we want update $\rho^{(t+1)}$ in such a way that the constraints are still met. Let

$$A = \begin{pmatrix} 0 & 1 & 1 & \cdots & 1 \\ 0 & 0 & 1 & \cdots & 1 \\ \cdots & \cdots & \cdots & \cdots & \cdots \\ 0 & 0 & 0 & \cdots & 1 \end{pmatrix},$$

then the constraints can be expressed as

$$\begin{cases} A\rho^{(t+1)} \prec \mathbf{0} \\ \rho^{(t+1)} \prec \delta \end{cases} \Rightarrow \begin{cases} A\left(\rho^{(t)} + s\Delta\right) \prec \mathbf{0} \\ \rho^{(t)} + s\Delta \prec \delta \end{cases} \Rightarrow \begin{cases} A\rho^{(t)} + sA\Delta \prec \mathbf{0} \\ \rho^{(t)} + s\Delta \prec \delta \end{cases}. \quad (2.1)$$

First, we check whether $s = 1$ satisfy **??**. If the constraint is met, then set $s = 1$. Otherwise, we need to look for a proper $s$, let $\Delta_a = A\Delta$. Suppose that $\rho^{(t)}$ satisfies the constraint, we only need to check the positive element of $\Delta_a$ and $\Delta$. The problem reduces to the following problem.

Suppose we have

$$\begin{pmatrix} a_1 \\ \vdots \\ a_l \end{pmatrix} \prec \begin{pmatrix} c_1 \\ \vdots \\ c_l \end{pmatrix}$$

and

$$\begin{pmatrix} b_1 \\ \vdots \\ b_l \end{pmatrix} \succ \mathbf{0},$$

find a value $s > 0$ such that

$$\begin{pmatrix} a_1 \\ \vdots \\ a_l \end{pmatrix} + s \begin{pmatrix} b_1 \\ \vdots \\ b_l \end{pmatrix} \prec \begin{pmatrix} c_1 \\ \vdots \\ c_l \end{pmatrix}.$$

Here we find a solution that $s = t \min_j \left(\frac{c_j - a_j}{b_j}\right)$, $0 < t < 1$, because we have

$$a_i + sb_i = a_i + t \min_j \left(\frac{c_j - a_j}{b_j}\right) b_i$$

$$
\begin{aligned}
&\leq a_i + tb_i \frac{c_i - a_i}{b_i} \\
&= (1-t)\, a_i + tc_i \\
&< (1-t)\, c_i + tc_i = c_i.
\end{aligned}
$$

For computation we just set $t = 0.999$. By this method we can find solution $s_1$ and $s_2$ that $A\rho^{(t)} + s_1\Delta_a \prec \mathbf{0}$ and $\rho^{(t)} + s_2\Delta \prec \delta$. Then we obtain $s = \min\{s_1, s_2\}$.

To deal with the constraint that $\sum_{i=1}^{n} \rho_i = 0$ one more penalty $\frac{1}{2}\gamma\left(\mathbf{1}^{'}\rho\right)^2$ was added to the goal function with gradient $\gamma\mathbf{1}^{'}\rho$ and Hessian $\gamma\mathbf{1}\mathbf{1}^{'}$, which are added to the Newton update. We set $\gamma = 10^5$.
